# Modeling SARS-CoV-2 and Influenza Infections and Antiviral Treatments in Human Lung Epithelial Tissue Equivalents

**DOI:** 10.1101/2021.05.11.443693

**Authors:** Hoda Zarkoob, Anna Allué-Guardia, Yu-Chi Chen, Olive Jung, Andreu Garcia-Vilanova, Min Jae Song, Jun-Gyu Park, Fatai Oladunni, Jesse Miller, Yen-Ting Tung, Ivan Kosik, David Schultz, Jonathan Yewdell, Jordi B. Torrelles, Luis Martinez-Sobrido, Sara Cherry, Marc Ferrer, Emily M. Lee

## Abstract

Severe acute respiratory syndrome coronavirus-2 (SARS-CoV-2) is the third coronavirus in less than 20 years to spillover from an animal reservoir and cause severe disease in humans. High impact respiratory viruses such as pathogenic beta-coronaviruses and influenza viruses, as well as other emerging respiratory viruses, pose an ongoing global health threat to humans. There is a critical need for physiologically relevant, robust and ready to use, *in vitro* cellular assay platforms to rapidly model the infectivity of emerging respiratory viruses and discover and develop new antiviral treatments. Here, we validate *in vitro* human alveolar and tracheobronchial tissue equivalents and assess their usefulness as *in vitro* assay platforms in the context of live SARS-CoV-2 and influenza A virus infections. We establish the cellular complexity of two distinct tracheobronchial and alveolar epithelial air liquid interface (ALI) tissue models, describe SARS-CoV-2 and influenza virus infectivity rates and patterns in these ALI tissues, the viral-induced cytokine production as it relates to tissue-specific disease, and demonstrate the pharmacologically validity of these lung epithelium models as antiviral drug screening assay platforms.

## Introduction

Newly emerging viral pathogens such as the severe acute respiratory syndrome coronavirus 2 (SARS-CoV-2) and other re-emerging respiratory viral threats, including influenza viruses, are a constant burden to human public health. A year after the initial discovery of the novel SARS-CoV-2 in 2019 [1, 2], the coronavirus disease 2019 (COVID-19) pandemic remains largely uncontrolled. While global vaccination efforts are underway, they are challenged by the emergence of viral variants of concern (VoC) that can potentially escape humoral immunity [3, 4, 6, 7]. Due to immune escape as well as the potential for newly emergent CoV strains, identifying effective anti-CoV drug is critical [8].

Several established cell lines permissive to SARS-CoV-2 infection *in vitro* have been used for high-throughput antiviral drug screening (HTS) efforts, including the African green monkey kidney Vero E6 cells, human hepatoma Huh7 and Huh7.5, colon carcinoma Caco2 cells, lung adenocarcinoma Calu-3, and angiotensin converting enzyme 2 (hACE2) overexpressing adenocarcinoma A549 cells, HEK293T cells, and several other non-human cell lines [4, 9-16]. While these cell lines are important tools for viral research, there is now evidence that many of the antiviral activities discovered are limited to the cells used for screening. For example, while hydroxychloroquine potently blocks SARS-CoV-2 infection of Vero E6 cells and Huh7 [4, 17], it is inactive against human Calu-3 lung cells [10, 18], and in both prophylactic and post-infection SARS-CoV-2’s in either rhesus macaque or golden Syrian hamster models [19-21]. Hydroxychloroquine is also ineffective in randomized COVID-19 human clinical trials [22, 23]. Animal models have been developed for pre-clinical drug development of COVID-19, but the low-throughput nature of high biocontainment models limits drug screening. There remains therefore a critical need for *in vitro* pre-clinical assays that are highly predictive of clinical drug efficacy, which can be used to prioritize compounds selection for animal testing.

Air-liquid interface (ALI) lung cultures provide a bridge between cultured cell lines and animal models [24-30]. In addition to more closely replicating the physiological environment of the human lung epithelium, ALI lung cultures support the replication of human coronaviruses (HCoV) with limited host cell range including HCoV-229E, HCoV-HKU1, HCoV-NL63, and HCoV-OC43 [29, 31-34]. To address the current shortcomings of traditional antiviral screening models, we investigated the use and predictive efficacy of ALI lung tissue equivalents modeling two regions of the lower respiratory tract – the tracheobronchial region and the alveolar region – in the context of SARS-CoV-2 and influenza virus infections.

In light of the critical role for cytokines in severe COVID-19 development, we further investigated the tissue-specific cytokine profiles of both tissue equivalents in response to SARS-CoV-2 compared to influenza infection. Cytokines likely play a multifaceted role in SARS-CoV-2 infection. On one hand, appropriate cytokine and chemokine induction is important for local viral clearance in tissues, whereas excessive cytokine production and altered cytokine balance is a key component of the COVID-19-associated cytokine storm [35-41]. While elevated serum cytokines associated with severe COVID-19 have been reported, major inflammatory contributing cytokines in locally infected lung tissue remain unknown [41, 42], in part because of the difficulty to measure intratissue cytokine levels in live patients during the course of infection and the relatively short half-life of secreted cytokines. The ability to model cytokine production in normal human tissues *in vitro* may help define the local *vs*. systemic cytokine production profiles.

We established the cellular complexity of lung epithelial ALI tissue models for both SARS-CoV-2 and influenza, their infectivity rates and patterns, host cytokine production as it relates to viral infection progression to disease, and demonstrate the pharmacologically validity of these *in vitro* models as antiviral drug screening platforms using viral protein immunostaining fluorescence imaging assays.

## Results

### Human tracheobronchial and alveolar ALI lung tissue equivalents present lung epithelial cell differentiation and expression of known SARS-CoV-2 host entry co-factors

ALI alveolar tissue equivalents are comprised of lung epithelial alveolar type I and type II cells, pulmonary fibroblasts and endothelial cells. In contrast, the tracheobronchial tissue equivalents are comprised of ciliated cells, goblet or secretory cells, and basal cells. hACE2 is a known host entry factor for both SARS-CoV and SARS-CoV-2 [43, 44] that is highly expressed (as determined by scRNAseq studies) in alveolar type II cells (ATII) and also expressed at lower levels in ciliated cells and some goblet cells [45-47]. By immunofluorescence with well characerrized antibodies (antibodies listed in Supplementary Table 1) we first examined expression of ACE2 and established lung cell marker antigens in both human alveolar and tracheobronchial ALI cultures. ACE2 was robustly expressed in the apical epithelium in both post-day 21 tracheobronchial ALI cultures **(Fig. 1a, top row**), and apical epithelium of post-day 21 alveolar ALI cultures **(Fig. 1b, top row)**. Type II transmembrane serine protease TMPRSS2 is another well-characterized virus entry host co-factor for both SARS-CoV-2, SARS-CoV, MERS-CoV, and influenza viruses, and transmembrane glycoprotein neuropilin-1 (NRP-1) enhances furin-dependent SARS-CoV-2 infection *in vitro* [44, 48-52]. We also observed robust TMPRSS2 and NRP-1 expression in both tracheobronchial and alveolar tissues using antibody-mediated detection (**Fig. 1a, 1b, second and third rows**). As expected, tracheobronchial tissue equivalents also exhibited robust staining acetylated α-tubulin as a marker for ciliated cells (α-tubulin+), mucin 5AC which is expressed on goblet or secretory cells (MUC5AC+) [53], and cytoskeletal protein cytokeratin 5 expressed in basal cells (KRT5+) [54], whereas alveolar tissue equivalents stained positive for actin filaments (Phalloidin+), alveolar type I (AQP5+, not shown) and alveolar type II (SP-B+) markers (**Figs. 1** and **2**). In tracheobronchial tissues, we observed positive co-staining of either ACE2, TMPRSS2, or NRP-1 with both α-tubulin+ ciliated cells as well as MUC5AC+ goblet cells (**Fig. 1a**). While most ACE2, TMPRSS2, or NRP-1 co-expressed with SP-B positive cells in alveolar tissues, a small minority of cells that were SP-B negative also expressed the probed SARS-CoV-2 entry cofactors **(Fig. 1b)**. Flow cytometry analysis of dissociated tissue cultures to identify major cell types are shown in **Supplemental Figure 1**.

**Figure 1:**
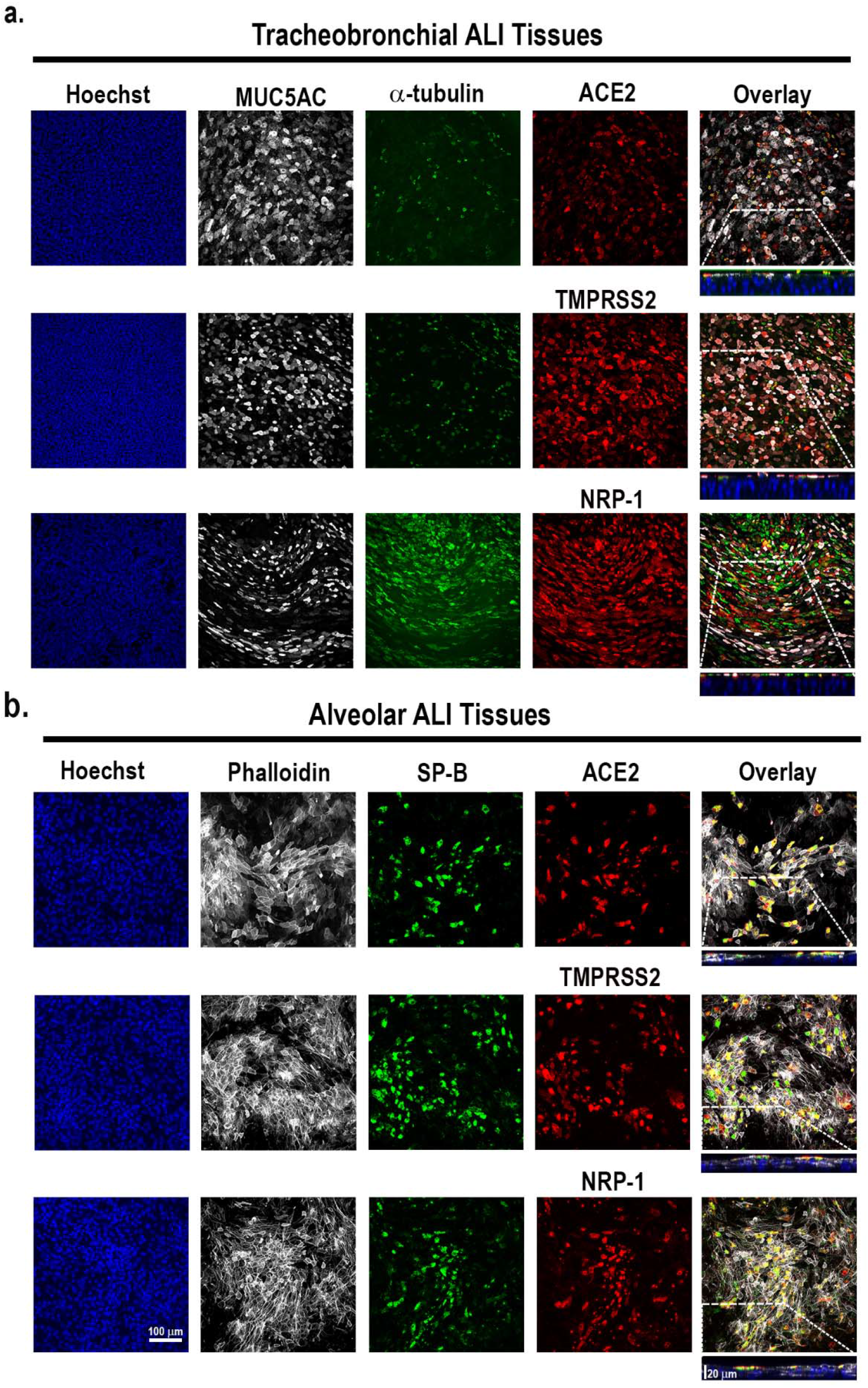
Apical expression patterns of known SARS-CoV-2 entry co-factors in tracheobronchial and alveolar ALI tissue equivalents. Post-day 21 tissues were stained with antibodies targeting hACE2, TMPRSS2, and NRP-1, as well as tissue-specific markers. a) Representative stained images of tracheobronchial ALI tissues with Hoechst (nuclei marker, blue), α -tubulin (ciliated cell marker, green), MUC5B (goblet cell marker, white) and hACE2 (top panel, red), TMPRSS2 (middle panel, red) or NRP-1 (bottom panel, red). The overlay image represents the maximum intensity projection of stained markers. The y/z plane cross section taken from the highlighted portion shows the selective apical expression of hACE2, TMPRSS2, and NRP-1. **b)** Representative stained images of alveolar ALI tissues with Hoechst (nuclei marker, blue), surfactant protein B (SP-B, ATII/pneumonocyte type II cell marker, green) and phalloidin (f-actin, white) co-stained with hACE2 (top panel, red), TMPRSS2 (middle panel, red) or NRP-1 (bottom panel, red). The overlay image represents the maximum intensity projection of stained markers and a y/z plane cross section from the highlighted portion shows the selective apical expression of hACE2, TMPRSS2, or NRP-1 in the tissues, contrasted to phalloidin, which is present throughout the tissue cross-section. Scale bar is 100 μm. Cross-section scale bar is 20 μm.

**Figure 2:**
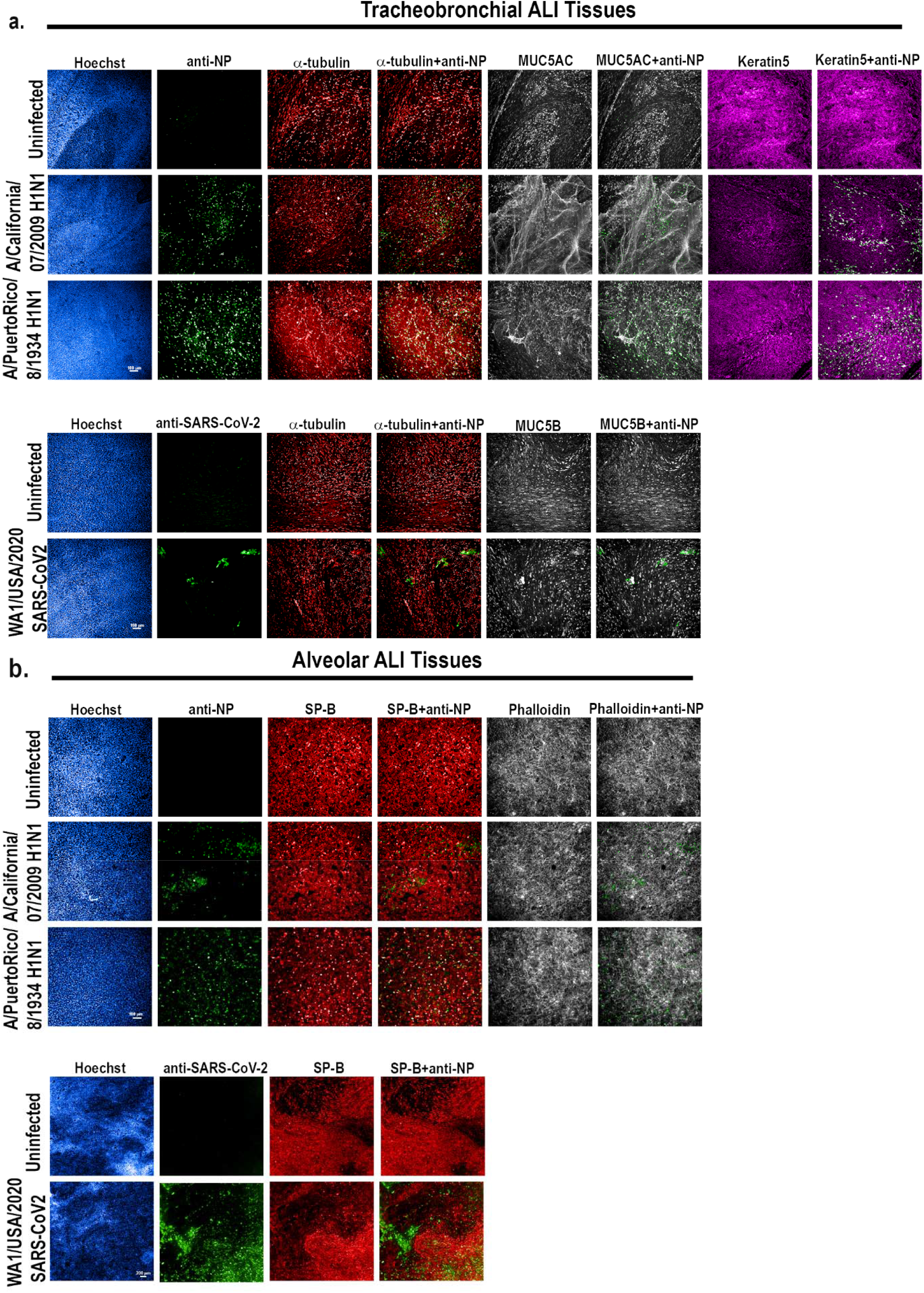
IAV and SARS-CoV-2 productively infect tracheobronchial and alveolar ALI tissue equivalents. Tracheobronchial and alveolar ALI tissues were infected with IAV strains pH1N1 or PR8 (1×10e5 TCID50 units), or SARS-CoV-2 (1e5 TCID50 units), (n=3). Infected tissues were fixed for 24 hpi for IAV inoculated tissue, 36 hpi for SARS-CoV-2 inoculated tracheobronchial ALI tissue, or 144 hpi for SARS-CoV-2 inoculated alveolar ALI tissue and stained with antibodies against selected cell markers and virus antigens as indicated: **a)** Tracheobronchial ALI tissues were stained with anti-α-tubulin (ciliated cell marker, red), anti-MUC5AC or MUC5B (goblet cell markers, white), and anti-keratin 5 (basal cell marker, magenta), along with anti-IAV N protein (green, top three panels) or anti-SARS-CoV-2 (monoclonal antibody cocktail targeting S and N proteins, green, bottom two panels) as the marker of infected cells. **b)** Alveolar tissues were stained with anti-surfactant protein B (SP-B, ATII cell marker, red), phalloidin (F-actin, general cell marker, white) along with anti-IAV N protein (green top three panels) or anti-SARS-CoV-2 (green, bottom two panels) as the marker of infected cells. Scale bar is 100 μm and 200 μm in IAV and SARS-CoV-2 infected tissues, respectively.

### SARS-CoV-2 and IAV productively infect human tracheobronchial and alveolar ALI tissue equivalents

We next determined whether SARS-CoV-2 and/or influenza A viruses (IAV) can infect human alveolar and tracheobronchial tissue equivalents. For this, we infected both tissues with either H1N1 IAV strains A/Puerto Rico/8/1934 (PR8) or pandemic A/California/07/2009 (pH1N1), or the clinical isolate SARS-CoV-2/WA/2020/D614, at an approximate multiplicity of infection (MOI) of total cells of 0.1-0.2. At indicated time-points (24 hpi for IAV and 36 hpi for SARS-CoV-2) after infection, we inactivated the virus by whole tissue fixation and stained for SARS-CoV-2 or IAV viral antigens, cell-type specific markers, and F-actin to observe tissue integrity, infection rates, and infected cell types **(Supplementary Table 1)**. In IAV exposed tracheobronchial tissue equivalents, we observed infection of multiple cell types at 24 hpi, with basal cells being the dominant infected cell type, followed by ciliated cells and goblet cells (**Fig. 2a, Supplemental Fig. 2a and 2b**). In SARS-CoV-2 infected tracheobronchial tissue equivalents, we observed co-staining of SARS-CoV-2 N antigen with both ciliated and goblet cells 36 h after initial infection **(Fig. 2a; Supplemental Fig. 2c)**, in agreement with the location of known SARS-CoV-2 host entry co-factors, as described above. In alveolar tissues, the majority of cells infected by IAV or SARS-CoV-2 were positive for SP-B, indicating primary infection of AT-IIs, although another SP-B negative cell subpopulation was also positive for SARS-CoV-2 N antigen **(Fig. 2b)**.

### Human tracheobronchial and alveolar ALI tissue equivalents exhibit slower viral infection kinetics for SARS-CoV-2 compared to IAV

Unlike IAV, which has a relatively short clinical incubation period of 1.5-2 days, SARS-CoV-2 has an average incubation period of 5.5 days after initial infection [55]. Thus, we investigated whether this is reflected in the tracheobronchial and alveolar lung tissue equivalents. Our data indicate that the number of IAV PR8 infected cells (MOI 0.1, per total cells) in both tracheobronchial and alveolar tissues rapidly spiked at 24 hpi, followed by a gradual decline in both infected ALI systems (**Fig. 3a** and **b**) and in secreted virus (**Fig. 3c**), in agreement with the robust and rapid replication described in the human host. Infection with IAV pH1N1 (MOI 0.1) resulted in the highest staining of IAV antigen at 48 hpi in both ALI tissue systems (**Fig. 3a** and **3b**), with peak viral release at 48 hpi in alveolar ALI tissues and 72 hpi in tracheobronchial ALI tissues, followed by a decline in infectious virion production by 144 hpi post-infection (**Fig. 3c**). Conversely, SARS-CoV-2 replication in both tissues was slower and of less magnitude (as shown by the total viral titers/insert shown in **Fig. 3c**) when compared to IAV, and in agreement with a longer incubation period in the human host. Indeed, the number of SARS-CoV-2 N antigen positive cells in both tissues was low at early timepoints, but continued to increase steadily over time in infected tissues independently of the MOI used for the stainings (0.1 to 1), with tracheobronchial tissues exhibiting peak infection at 72 hpi (**Fig. 3a**) and alveolar tissues at the latest time-point tested (144 hpi) (**Fig. 3b**). Correspondingly, infectious virion production from SARS-CoV-2 infected alveolar tissues peaked at 144 hpi indicating a steady and increasing infection over the tested 6 days period (**Fig. 3c**). In tracheobronchial tissues, viral production was also dose-dependent, although infection with higher MOIs (approximately 3 and 10) resulted in a higher virion production at 24 hpi, followed by a decline in later timepoints, which may be due to cell death as a result of an initially high viral exposure (**Fig. 3c**). Interestingly, alveolar tissues seemed to be more susceptible to IAV and SARS-CoV-2 infection than tracheobronchial cultures, as evidenced by the higher number of infected cells (**Fig. 3b**) and viral titers (**Fig.3c**). Cellular markers for each time-point are shown in **Supplemental Fig. 2**. Thus, both IAV and SARS-CoV-2 can productively infect lung tissue equivalents, though with different kinetics that may contribute to their different clinical incubation periods.

**Figure 3:**
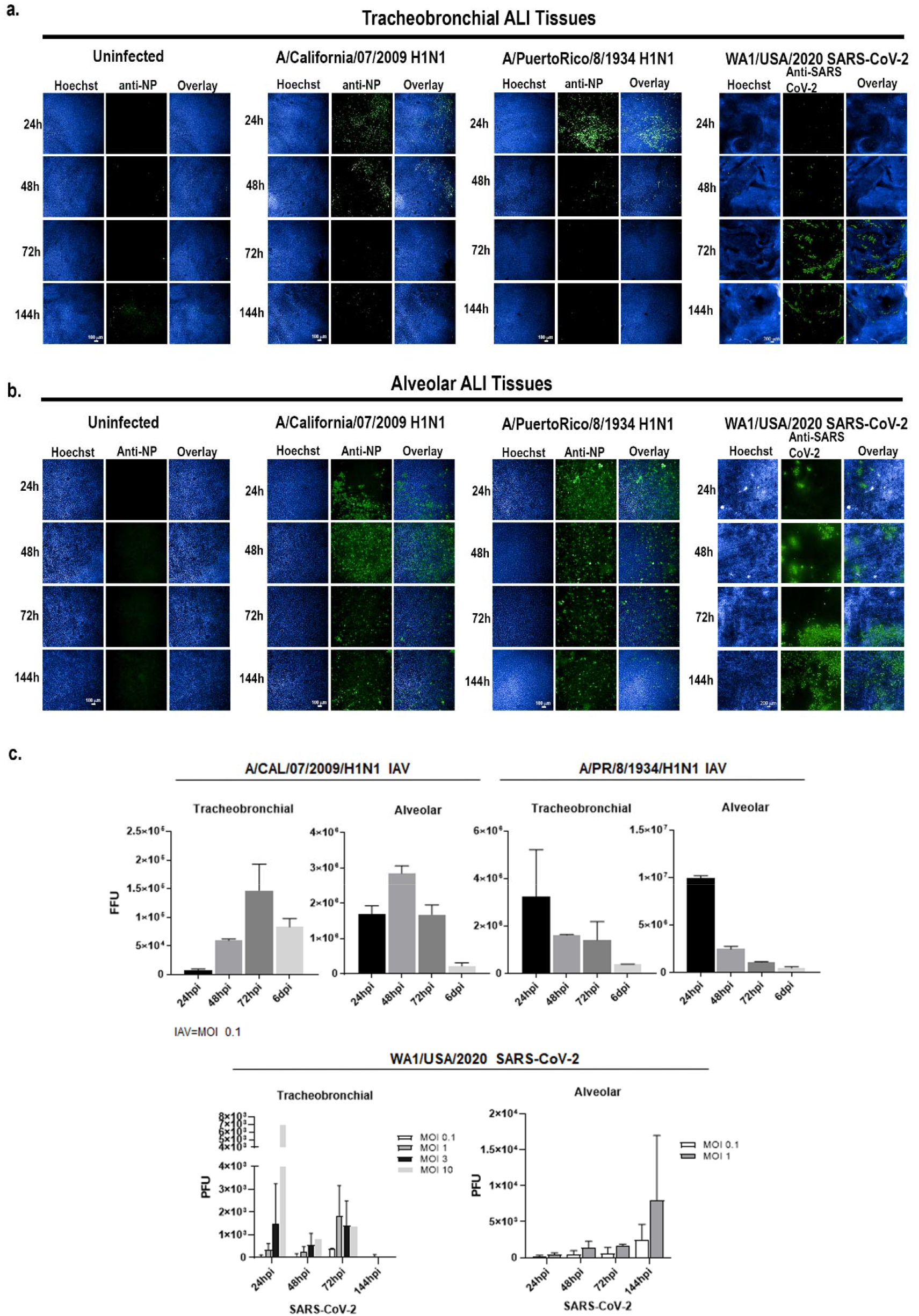
IAV and SARS-CoV-2 exhibit different infection kinetics in tracheobronchial and alveolar ALI tissue equivalents. Tracheobronchial and alveolar ALI tissues were infected with IAV pH1N1 or PR8 at approx. MOI of 0.1, and SARS-CoV-2 at MOI of 1 (fixed tissue samples shown) or as indicated in titer plots. Apical washes were collected and tissues fixed at 24, 48, 72 and 144 hpi. **a)** Tracheobronchial and **b)** alveolar ALI tissues were stained with anti-IAV N protein and anti-SARS-CoV-2 to label infected cells (shown in green) as well as the nuclear dye Hoechst (blue). c) Production of infectious virus from the apical chamber of tracheobronchial or alveolar ALI tissues after exposure to pH1N1 (MOI of 0.1), PR8 (MOI of 0.1), or SARS-CoV-2 (MOI of 0.1 and 1 for alveolar tissues; MOIs of 0.1, 1, 3, and 10 for tracheobronchial tissues) at 24, 48, 72, or 144 hpi. IAV titers were measured using a focus forming unit assay on LLC-MMK2 SIAT1 cells, SARS-CoV-2 was measured using plaque assay on Vero E6 cells, and expressed as total FFU (IAV) or PFU (SARS-CoV-2)/tissue. Data is represented as M±SD for a minimum of n=2 independent experiments/biological replicates. No virus was detected in uninfected controls (data not shown).

### Human tracheobronchial and alveolar ALI tissue equivalents exhibit distinct inflammatory cytokine profiles in response to IAV or SARS-CoV-2 infection

Secretion of pro-inflammatory cytokines during respiratory viral infection by SARS-CoV-2 or IAV is a major contributor to severe COVID-19 disease and severe influenza in humans, respectively [1, 56-59]. The cellular sources of these cytokines and underlying inflammatory mechanisms, however, remain uncertain. It has been proposed that both lung epithelial cells and alveolar macrophages may directly produce local tissue cytokines and immune markers in response to SARS-CoV-2 infection [45, 60, 61]. In IAV infection, the interplay between epithelial, endothelial, and innate immune cells is likely critical to cytokine storm development, with pulmonary endothelium playing a critical amplifier role[59, 62]. Although our lung tissue equivalents modeled in this study lack the myeloid cellular compartment, which plays an important role in the innate immune response, we were still able to investigate alveolar with endothelium *vs*. tracheobronchial epithelial chemokine and cytokine production in response to viral infection. To determine this tissue-specific immune response, we infected both lung tissue equivalents at indicated MOIs and sampled basal media supernatants at 24, 48, 72, and 144 hpi. We used a a custom multiplex Luminex assay to measure COVID-19 associated cytokines/chemokines [1, 56, 57, 63] including interleukin 6 (IL-6), C-X-C motif chemokine ligand 10 or interferon gamma-induced protein 10 (CXCL-10/IP-10), C-C motif ligand 3 or macrophage inflammatory protein 1 alpha (CCL3/MIP-1α), C-C motif ligand 2 or monocyte chemoattractant protein-1 (CCL2/MCP-1), interleukin-1 receptor antagonist 1a (IL-1RA) and granulocyte colony-stimulating factor (G-CSF), as well as several other Th1/Th2/Th17 cytokines and interferons **(Fig. 4, Fig. 5, Supplemental Fig. 3)**.

**Figure 4:**
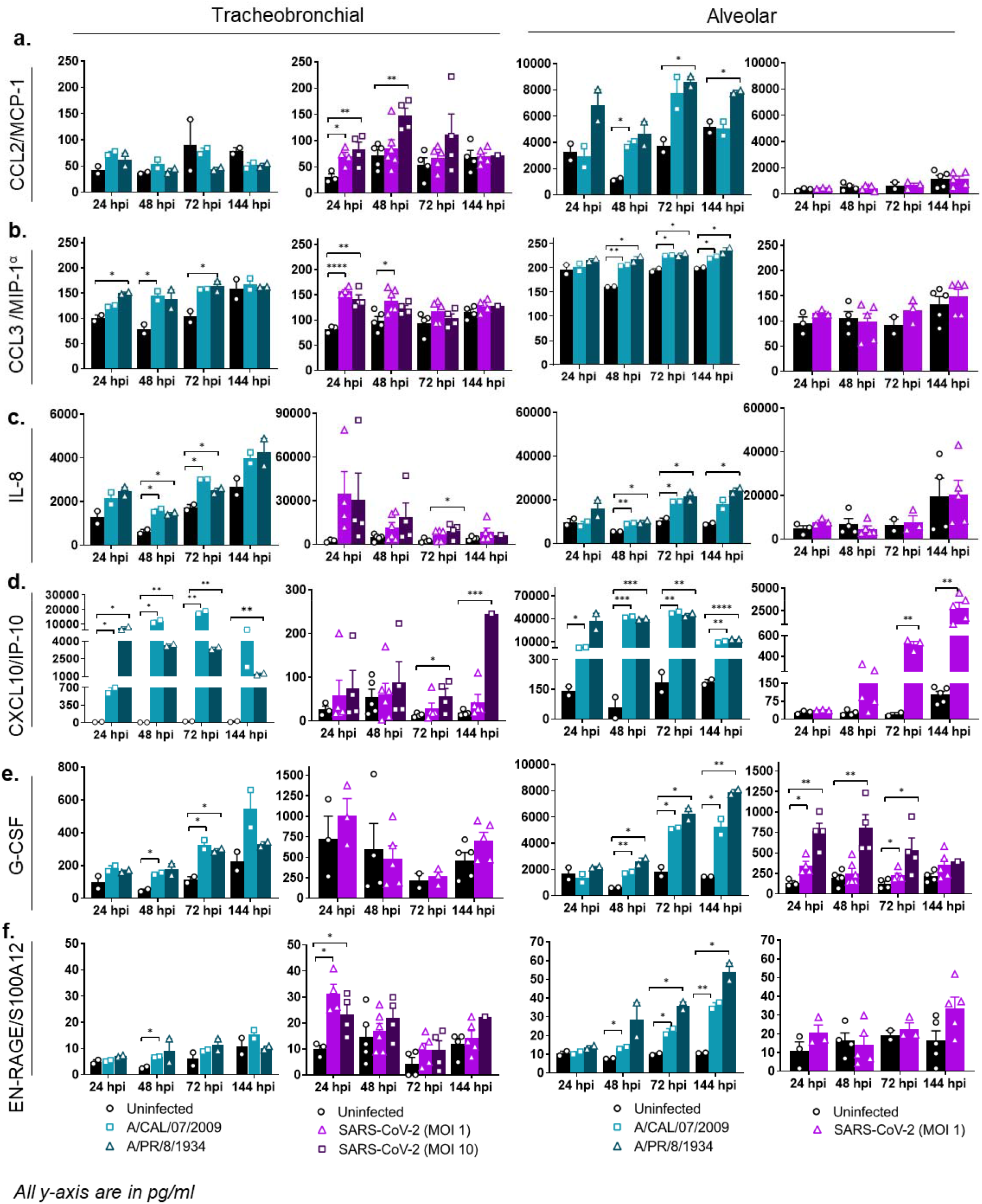
Alveolar and tracheobronchial ALI tissues produce tissue-specific chemokines and growth factors in response to IAV and SARS-CoV-2 infection. Basal compartment media were collected from tracheobronchial (left two panels) or alveolar (right two panels) ALI tissues at indicated time-points and analyzed for cytokine and chemokine secretion by Luminex assay. IAV infected tissues (approx. MOI of 0.1) are represented in shades of teal, where light teal shows infection with the IAV pH1N1 strain and dark teal shows infection with the IAV PR8 strain, whereas SARS-CoV-2 infected tissues are represented in shades of purple, with progressing color from low MOI (1) to high MOI (10): (**a**) CCL2/MCP-1, (**b**) CCL3/MIP-1α, (**c**) IL-8, (**d**) CXCL10/IP-10, (**e**) G-CSF, (**f**) EN-RAGE/S100A. All measurements on y axis are in pg/ml. Data is represented as M±SEM for a minimum of n=2 independent experiments and/or biological replicates; Student t-test of IAV or SARS-CoV-2 infected tissues *vs*. uninfected controls at each timepoint: \**p*⍰<⍰0.05, \*\**p*⍰<⍰0.005, \*\*\**p*⍰<⍰0.0005, \*\*\*\**p*⍰<⍰0.00005.

**Figure 5:**
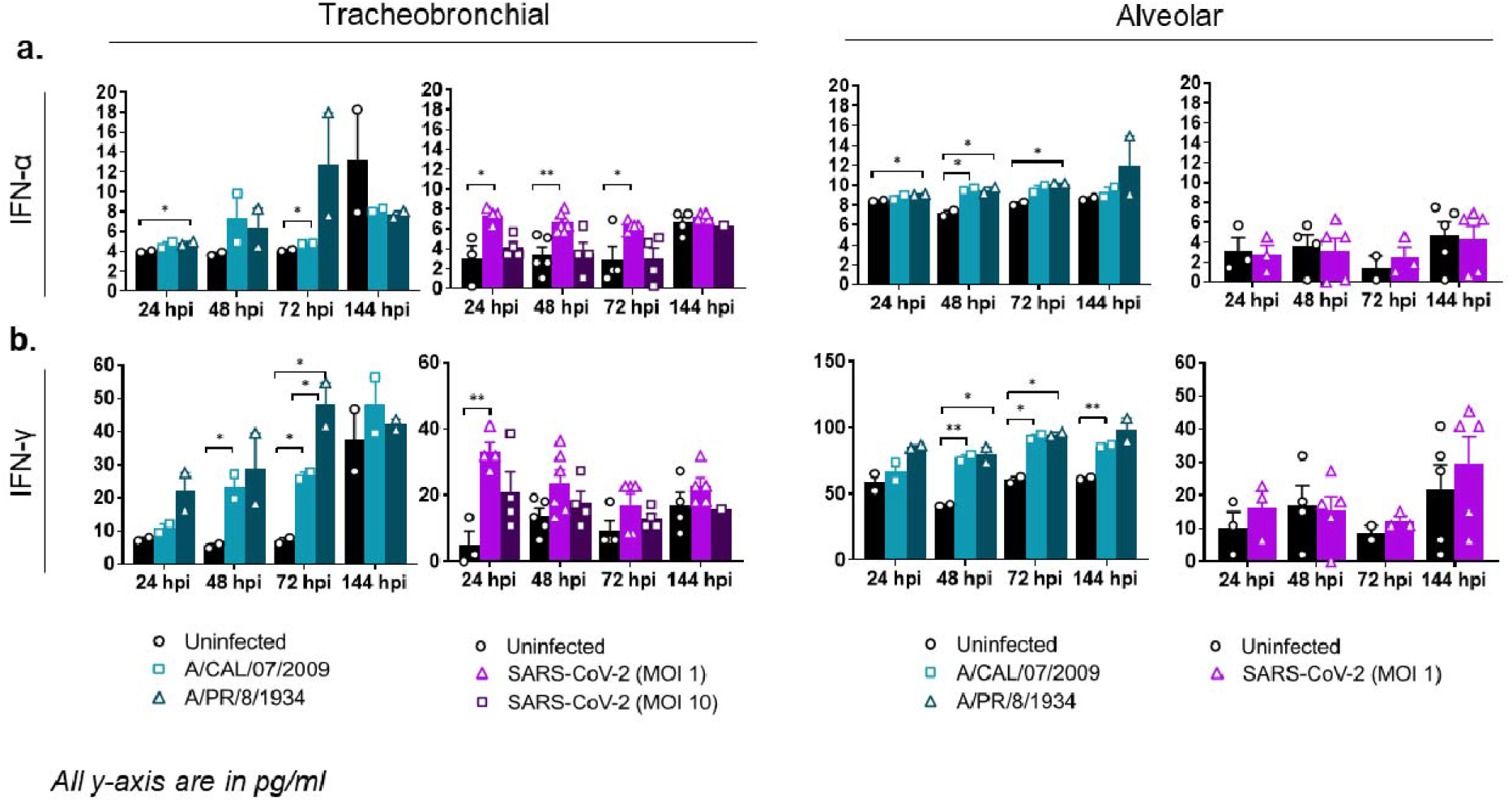
Alveolar and tracheobronchial ALI tissues produce moderate levels of IFN in response to IAV and SARS-CoV-2 infection. Basal compartment media were collected from tracheobronchial (left two panels) or alveolar (right two panels) ALI tissues at indicated time-points and analyzed for cytokine and chemokine secretion by Luminex assay. IAV infected tissues (approx. MOI=0.1) are represented in shades of teal, where light teal shows infection with the IAV pH1N1 strain and dark teal shows infection with the IAV PR8 strain, whereas SARS-CoV-2 infected tissues are represented in shades of purple, with progressing color from low MOI (1) to high MOI (10): (**a)** IFN-α, (**b)** IFN-γ. All measurements on y axis are in pg/ml. Data is represented as M±SEM for a minimum of n=2 independent experiments and/or biological replicates; Student t-test of IAV or SARS-CoV-2 infected tissues *vs*. uninfected controls at each timepoint: \**p*⍰<⍰0.05, \*\**p*⍰<⍰0.005, \*\*\**p*⍰<⍰0.0005, \*\*\*\**p*⍰<⍰0.00005.

We observed that the level of induction for most of the immune markers varied significantly in a virus- and tissue-dependent manner (**Fig. 4, Fig. 5, Supplemental Fig. 3**). Overall, tissues infected with influenza virus showed a stronger immune response **(Fig. 4, Fig. 5)**, corresponding with the higher viral titers reported in comparison to SARS-CoV-2 infection **(Fig. 3)**. Chemokine levels were significantly increased in both tracheobronchial and alveolar tissues infected with either A/Cal/07/2009/H1N1 or A/PR/8/1934/H1N1 influenza strains (with the exception of CCL2/MCP-1 in tracheobronchial tissues), showing an early and robust response at 24 or 48 hpi, correlated with maximum viral production **(Fig. 3)**, that was maintained until later timepoints **(Fig. 4)**. Interestingly, chemokine production after SARS-CoV-2 infection was greatly dependent on tissue type and appeared to be mostly restricted to tracheobronchial tissues, which showed significantly higher production compared to uninfected controls at either early (CCL2/MCP-1 and CCL3/MIP-1α, **Fig. 4a, Fig. 4b**) or later (CXCL-8/IL-8 and CXCL-10/IP-10, **Fig. 4c, Fig. 4d**) timepoints. CXCL-10/IP-10, an important chemoattractant for neutrophils, was the only chemokine in our panel induced in SARS-CoV-2 infected alveolar tissues, starting at 48 hpi and significantly increased at 72 and 144 hpi (**Fig. 4d**), corresponding to the increased number of infectious viral particles at later timepoints (**Fig. 3c**). The increased chemokine levels observed in lung tissue equivalents, especially in the tracheobronchial model, are in agreement with the chemokine storm observed in severe human COVID-19 cases and different SARS-CoV-2 experimental models [64], where the role of CXCL-10 has been particularly highlighted in COVID-19-associated Acute Respiratory Distress Syndrome (ARDS) as well as in severe influenza[59].

We next examined inflammatory Th1, Th2, and Th17 immune responses to IAV and SARS-CoV-2 infection, including type I (IFN-α and IFN-β), type II (IFN-γ), and type III (IFN-λ) interferons. Although not in high amounts, we observed a modest but significant type I/II interferon response in some of the IAV and SARS-CoV-2 infected tissues compared to the uninfected controls (**Fig. 5**). IL-28A/IFN-λ2 protein levels were below the limit of detection. Specifically, in SARS-CoV-2 infected tissues, IFN-α and IFN-γ levels were increased in the tracheobronchial model but not in the alveolar, suggesting a tissue-specific response to the viral infection at early timepoints, whereas IFN-β was not detected in either tissue (**Fig. 5, Supplemental Fig**.**3**). In addition, most Th1 (TNF-α, TNF-β, IL-2), Th2 (IL-1β, IL-6, IL-10, IL-18), and Th17 (IL-17) immune markers tested were elevated in both IAV infected tissue equivalents compared to uninfected controls at several timepoints (**Fig. 6, Supplemental Fig. 3**), demonstrating a modest but sustained inflammatory response to influenza, especially in the alveolar model. In contrast, the cytokine response to SARS-CoV-2 infection was again mostly observed in tracheobronchial cultures, with highest detected secretion at early timepoints but decreasing after 72 hpi (with the exception of IL-10 that was maintained until 144 hpi, **Fig. 6d**) (Fig. 6, **Supplemental Fig. 3**). Only TNF-α (Fig. 6a), IL-10 (Fig.6d), and IL-18 (**Supplemental Fig. 3**) showed moderate but significant increased levels in alveolar tissues at 72 and 144 hpi, respectively, in correlation with maximum viral production (**Fig. 3**). IL-1β was not detected in either tissue during SARS-CoV-2 infection.

**Figure 6.**
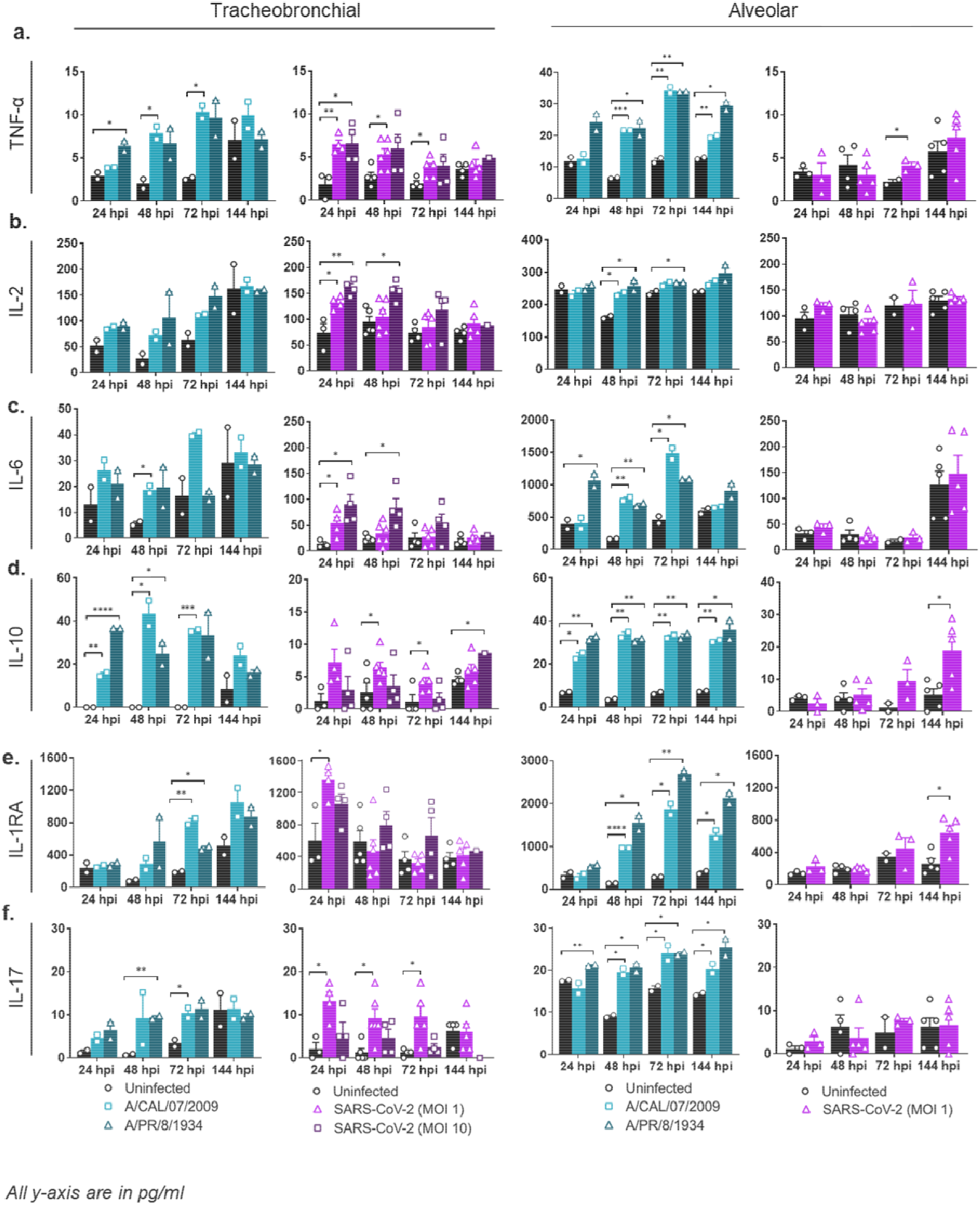
Production of Th1/Th2/Th17 markers. Basal compartment media were collected from tracheobronchial (left two panels) or alveolar (right two panels) ALI tissues at indicated time-points and analyzed for cytokine and chemokine secretion by Luminex assay. IAV infected tissues (MOI of 0.1) are represented in shades of teal, where light teal shows infection with the IAV pH1N1 strain and dark teal shows infection with the IAV PR8 strain, whereas SARS-CoV-2 infected tissues are represented in shades of purple, with progressing color from low MOI (1) to high MOI (10): (**a**) TNF-α, (**b**) IL-2, (**c**) IL-6, (**d**) IL-10, (**e**) IL-1RA, (f) IL-17. All measurements on y axis are in pg/ml. Data is represented as M±SEM for a minimum of n=2 independent experiments and/or biological replicates; Student t-test of IAV or SARS-CoV-2 infected tissues *vs*. uninfected controls at each timepoint: \**p*⍰<⍰0.05, \*\**p*⍰<⍰0.005, \*\*\**p*⍰<⍰0.0005, \*\*\*\**p*⍰<⍰0.00005.

We also measured secreted IL-6, which has been associated with the acute inflammatory response and cytokine storm characteristic of severe COVID-19 and severe influenza, as well as the increased production of anti-inflammatory cytokine IL-10 to maintain immune homeostasis [57, 65-67]. The presence and ratio of IL-6 and IL-10 may be used as predictors of COVID-19 disease severity. In our SARS-CoV-2 infected tissue lung equivalents, IL-6 was only increased in tracheobronchial cultures at 24 and 48 hpi (slightly decreasing at 72 hpi and normalizing to uninfected control levels at 144 hpi), whereas IL-10 was significantly higher in both tracheobronchial and alveolar cultures starting at 48 hpi and maintained at later timepoints (**Fig. 6c, Supplemental Fig. 3**). In addition, both IAV and SARS-CoV-2 infected tissues showed an elevated production of another anti-inflammatory protein, IL-1RA (**Fig. 6e**), indicating that these models are capable of producing reported COVID-19 associated molecules to suppress inflammation during viral infection.

Lastly, we observed increased production of other immune markers such as G-CSF, which controls neutrophil development and function, and EN-RAGE/S100A12, which participates in the migration and recruitment of leukocytes and promotes cytokine and chemokine production, as well as IL-7 and hepatocyte growth factor (HGF) which can mediate T-cell development and are markers of neutrophil activation. All four markers have been found elevated in the serum of hospitalized COVID-19 patients [68, 69], and increased levels have also been correlated with the cytokine storm associated with COVID-19 disease severity. We observed increase of all four proteins in response to both IAV and SARS-CoV-2 in tracheobronchial tissues up to 72 hpi; but not in alveolar cultures infected with SARS-CoV-2 **(Fig. 4e, Fig. 4f, Fig. 6f, Supplemental Fig. 3)**.

### Validation of antiviral drugs in lung tissue equivalents

To evaluate the ability of these physiological relevant ALI tissues to measure antiviral drug activity, we tested a panel of antiviral drugs on either tracheobronchial or alveolar cultures for antiviral activity. We first tested remdesivir, which has been granted emergency use authorization by the US Food and Drug Administration (FDA) for hospitalized COVID-19 patients [70], and type I interferon (IFN-β) for anti-SARS-CoV-2 activity. As expected, both remdesivir and IFN-β robustly inhibited SARS-CoV-2 infection as seen by qRT-PCR and direct SARS-CoV-2 antigen staining in the tracheobronchial tissue model (**Fig. 7, Supplemental Fig. 4)**. Furthermore, we tested two other compounds: cyclosporine, previously identified in a high-throughput screen with anti-SARS-CoV-2 infection activity in both human hepatoma (Huh7.5 cells) and human lung adenocarcinoma cells (Calu-3) in a monolayer cell-based model [10], as well as hydroxychloroquine. Neither cyclosporine nor hydroxychloroquine reduced SARS-CoV-2 infection in the tracheobronchial lung model (10μM, **Fig. 7a**, bottom two rows, **Fig. 7c, Supplemental Fig. 5a, 5c**). Furthermore, hydroxychloroquine failed to reduce SARS-CoV-2 viral production in the alveolar model (10μM, **Supplemental Fig. 5d**). However, when we tested camostat, a TMPRSS2 inhibitor, and nelfinavir, an anti-retroviral, currently being tested in clinical trials [44, 71-74], both reduced SARS-CoV-2 infection in the tracheobronchial ALI tissue (**Fig. 7a, 7c, Supplemental Fig. 5a**, 10μM) and alveolar ALI tissue (Supplemental Fig. 5b, 10μM).

**Figure 7:**
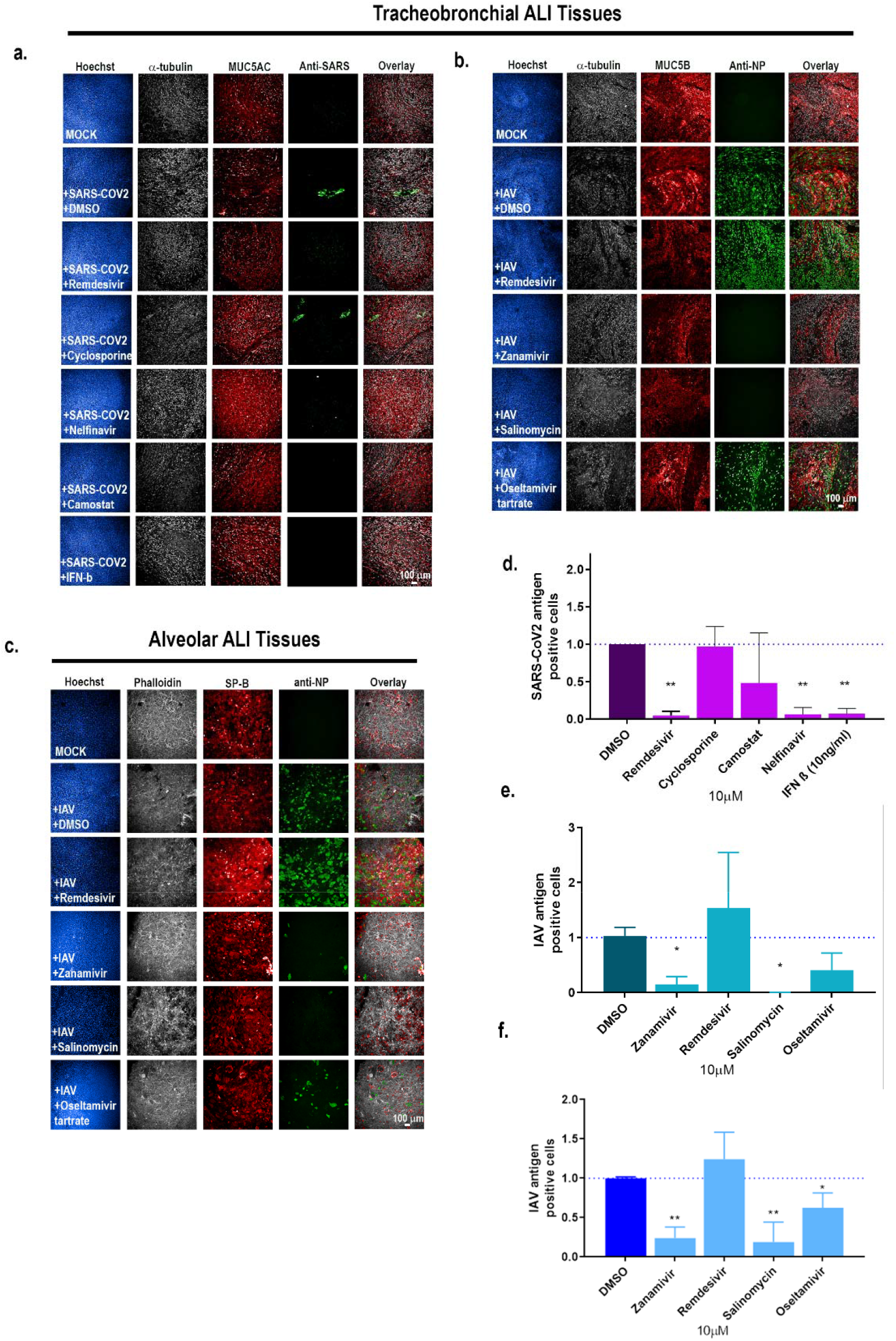
Tracheobronchial and alveolar ALI tissue equivalents predictively measure antiviral compound response in the context of SARS-CoV-2 and IAV infection. Selected compounds were added to the basal media chamber (10 μM final concentration) of the tracheobronchial and alveolar ALI tissues for 1 h and then infected with IAV PR8 (approx. MOI of 0.1), and SARS-CoV-2 (MOI of 0.1). IAV and SARS-CoV-2 infected tissues were fixed for 24 and 36 hpi, respectively, and stained with antibodies against selected cell markers and viral specific antigens. (**a, b)** Tracheobronchial ALI tissues were stained with anti-α-tubulin (ciliated cell marker, white), anti-MUC5AC or anti-MUC5B (goblet cell marker, red), along with anti-N protein (green, right five panels) and anti-SARS-CoV-2 (monoclonal antibody cocktail targeting S and N proteins, green) as the marker of infected cells. (**c**) Alveolar ALI tissues were stained with anti SP-B (ATII cell marker, red), phalloidin (F-actin, white), anti-N (green). Scale bar is 100 μm. (**d, e**) Image-based quantification of infected cells in compound treated and subsequent (**d**) SARS-CoV-2 or IAV infected tracheobronchial (**e**) or alveolar (**f**) ALI tissues. Data is represented as M±SD for a minimum of n=2 independent experiments and/or biological replicates; Student t-test of IAV or SARS-CoV-2 infected tissues vs. uninfected controls at each timepoint: *p⍰<⍰0.05, **p⍰<⍰0.005, ***p⍰<⍰0.0005, ****p⍰<⍰0.00005.

We next tested whether anti-SARS-CoV-2 compounds remdesivir and nelfinavir would also reduce IAV infection in the tracheobronchial (**Fig. 6b**) or alveolar (**Fig. 7c**) ALI tissue models. Neither remdesivir nor nelfinavir reduced IAV antigen staining in these tissues. However, clinically approved anti-IAV compounds zanamivir and oseltamivir, as well as the reported anti-IAV drug salinomycin, did inhibit IAV infection using a single dose approach in the tracheobronchial (**Fig. 7d**) ALI tissues at 10μM, as well as in a dose-dependent manner as expected (**Supplemental Fig. 5**). The TMPRSS2 inhibitor nafamostat also failed to block IAV infection in the tracheobronchial model, although we did observe a significant reduction in viral production (data not shown). Taken together, these findings validate the use of lung tissue equivalents for studying the efficacy of anti-viral drugs.

## Discussion

We have shown that both primary human tracheobronchial and alveolar ALI tissues in a transwell plate format are relevant models for studying multiple aspects of SARS-CoV-2 and IAV infection, in addition to being a valuable platform for antiviral drug testing. Progression of infection from the upper respiratory tract to the lower respiratory tract is necessary for both severe COVID-19 and influenza. Samples taken from patients with fatal COVID-19 have shown infection in ciliated bronchiolar epithelial cells, AT-IIs, goblet cells, club-like cells, and endothelial cells [75-77]. Similarly, in the described tissue models we see SARS-CoV-2 infection, peaking at 3-6 dpi, of both ciliated (α-tubulin+) and goblet (MUC5AC+) cell populations in the tracheobronchial ALI tissues, as well ATs in the alveolar ALI tissue model. We also observed robust SARS-CoV-2 and influenza infectious viral production in both tissues, although apical washes collected from alveolar tissues exhibited slightly higher SARS-CoV-2 titers than tracheobronchial (approx. 2×10^3^ PFU/tissue in apical wash of tracheobronchial ALI tissues and approx. 8×10^3^ PFU/tissue in an apical wash of alveolar ALI tissues). In contrast, a previously reported lung-on-chip model, consisting of primary human ATs and human lung microvascular endothelial cells, did not find productive infection with SARS-CoV-2 in ATs (Thacker et al., 2020). This may be due to the absence of hACE2 expression in ATs in the lung-on-chip model compared to robust hACE2 expression in the alveolar tissues in this study. While we did not observe SARS-CoV-2 N antigen in endothelial cells in our ALI tissue models, this may be because we only exposed the tissue apical side to the viral inoculum. In this regard, future studies will need to look at viral infection over an extended period to further characterize the full infection dynamics in the tracheobronchial and alveolar lung tissue equivalents, in addition to direct exposure of lung endothelial cells to SARS-CoV-2.

Elevated cytokine levels and pathogenic inflammation plays a central role in COVID-19 morbidity and mortality. Circulating chemokines, interferons, interleukins, growth factors and other pro-inflammatory cytokines as the main molecules involved in the development of the cytokine storm associated with COVID-19 severity, including IL-6, IL-2, IL-10, CXCL-10/IP-10, TNF, IFN-γ, CCL3/MIP-1α, CCL4/MIP1β, or G-CSF. In particular, high levels of IL-6 and TNF are strongly associated with increased mortality, and elevated levels of anti-inflammatory cytokines IL-10 and IL-1RA have also been correlated with disease severity and fatal outcome [1, 56, 57, 63, 65, 78]. Circulating cytokines observed in serum may not, however, represent local tissue cytokine levelswhich may be key potentiators of the systemic hyperinflammatory response. There have been extensive investigation into immunomodulatory drugs for the treatment of COVID-19 [42].

In agreement with previous COVID-19 reports, we did find a significant induction of IP-10/CXCL10 levels in both tracheobronchial and alveolar SARS-CoV-2 infected tissues (up to 25-fold increase in alveolar cultures compared to uninfected controls), as well as other important chemokines and growth factors (CCL2/MCP-1, CCL3/MIP-1α, CXCL-8/IL-8, G-CSF) that were elevated in tracheobronchial cultures. Interferon (IFN-α and IFN-γ) and Th1, Th2, and Th17 cytokine responses were only moderate but significant, and mostly restricted to tracheobronchial tissues. It included pro-inflammatory (IL-6 and TNF-α), and anti-inflammatory (IL-10 and IL-1RA) immune modulators, both which are known to have key roles in the pathogenesis associated to COVID-19 disease [1, 56, 57, 65, 78]. As demonstrated by the Luminex data, the cytokine induction after SARS-CoV-2 infection appeared to be tissue-dependent in our ALI models. Interestingly, even though alveolar tissues had a more robust infection compared to tracheobronchial cultures (as shown by the higher viral loads reported in **Fig. 3**), the cytokine and chemokine production was not as strong when compared to the response observed in tracheobronchial cultures. Still, both tissues had an overall immune response that corresponded with the infection dynamics, with alveolar tissues showing slight increases in some cytokines at later timepoints (72 to 144 hpi) compared to earlier responses (24 to 72 hpi) in the tracheobronchial model.

A major limitation of these models is that we were only able to study the local epithelial-driven response by itself, without the contribution of myeloid cells or lymphocytes, which are important in mounting an appropriate innate immune response against viral infections. In general, we observed that both tracheobronchial and alveolar ALI tissues are capable of mounting a local epithelial-driven response to IAV and SARS-CoV-2 virus infection. Overall, IAV infected tissues had a stronger and earlier inflammatory response than SARS-CoV-2 infected ALI tissues, including inflammatory as well as anti-inflammatory immune modulators. And although myeloid cells were not included in this model, we hypothesize that the addition of the myeloid compartment in both tissues will show a more pronounced immune response that may reflect more accurately the different stages of human COVID-19 and severe influenza disease progression. Future studies will address the contribution of the lung epithelium and its role in recruiting additional inflammatory immune cells.

While there has been intense high-throughput screening efforts to discover potential antivirals for SARS-CoV-2, relatively few compounds have proven to be effective in clinical settings. Most of the studies published have relied on traditional mono-cellular tissue culture models, which do not allow crosstalk among different cell populations. In some cases, relying on *in vitro* activity profiles may prove detrimental. This is the case for hydroxychloroquine, initially shown to have anti-SARS-CoV-2 activity *in vitro*, but later proven ineffective at reducing COVID-19 patient outcome or hospitalization stay [22, 23]. Interestingly, hydroxychloroquine failed to reduce SARS-CoV-2 infection in our *in vitro* ALI tissue equivalents, indicating that these 3D models might mimic better human tissue responses than other *in vitro* systems.

A 3D lung tissue model may be more high throughput and more accessible route for drug validation studies for newly emerged viral pathogens compared to small animal models, as they can readily be deployed and utilized in labs that do not specialize in animal models, do not require pharmacokinetics studies to be carried out prior to antiviral evaluation, and can easily be scaled up to include dose-response testing. While the study described here used 9mm and 12mm transwell inserts, these models can be readily adapted for use in a further miniaturized 96-well format which will increase throughput of the model for drug testing. In both the tracheobronchial and the alveolar ALI models, we were able to demonstrate robust antiviral activity of known anti-SARS-CoV-2 compound remdesivir, while also excluding hit compounds from 2D monolayer culture systems. Whether ALI tissue models can provide more accurate results regarding their efficacy *in vivo* requires to be further evaluated and must be taken into consideration with a compound’s pharmacokinetic properties. These models can also work in parallel with animal model efforts, by serving as an accessible and physiologically relevant human-based platform on which to prioritize compound selection for animal testing and further pre-clinical evaluation.

Several lung ALI models have been described within the last year to assess both IAV and CoV infectivity, including SARS-CoV-2, each with advantages and limitations. Small airway and alveolar lung-on-chip models have been published or reported in pre-prints [29, 79-83]. In addition, upper respiratory tract models such as nasal epithelium, which express high levels of SARS-CoV-2 receptors, have also been reported as successful *in vitro* culture systems for SARS-CoV-2 infection [84, 85]. Lung-on-chip models mostly recreate the interface between lung epithelial cells at the ALI and an endothelial cell monolayer. In many instances, the endothelial cells are exposed to liquid flow, and in some models, mechanical stretching is included in the chip design to mimic breathing[83]. Many of the lung-on-chip systems are low throughput, not readily compatible with laboratory automation used in drug screening facilities, and some cases chips are made of polydimethylsiloxane PDMS, thus limiting its use for drug testing. The use of a transwell-based multi-well plate assay platform as described in our study enables a versatile and modular approach for the future biofabrication of tissue ALI models with tailored physiological complexity and disease relevance. As an example, we have reported the use of bioprinting technique to create a vascularized skin tissue [86] and the same approach can be applied to recreate a vascularized lung ALI tissue model. Addition of non-lymphocyte immune cells has also been explored using transwell plates with biofabricated tissue equivalents [87] which can be applied to the lung ALI assay systems to assess the participation of innate immune cells in infectivity and COVID-19 relevant immune responses. Transwell multi-well plates are compatible for laboratory automation equipment for the implementation of drugs screens. Lung ALI tissue equivalents can be biofabricated in a 96-well transwell plate to enable a remarkable increase in the numbers of compounds that can be tested and therefore serve as an effective selection step for animal testing and reducing cost. We envision that newer lung ALI platforms will be developed based on these protocols and the benchmarks described in this study, and will include additional physiological features desired in a lung tissue model. In this regard, Zamprogno et al. have described a lung-on-chip platform that is open access and includes stretching, and it is high-throughput compatible[83].

In summary, we have described the characterization of two distinct lower respiratory tract lung epithelial ALI tissue models for studying SARS-CoV-2 and IAV infection in relation to complex tissue-related disease. We established differential cytokine profiles induced by tracheobronchial *vs*. alveolar tissues in response to two pandemic respiratory viruses, the recently emerged coronavirus SARS-CoV-2 and IAV, including a variant of the 2009 pH1N1. In addition, using known and novel antivirals we demonstrate the pharmacological validity of these two models as antiviral drug screening assay platforms. This characterization will serve as the foundation to biofabricate lung ALI tissue models, which may include additional physiological features that are relevant for the infection of respiratory viruses and the disease that they cause.

## Methods

### Viral Propagation

Vero E6 cells were obtained from the American Type Culture Collection (ATCC CRL-1586) and cultured at 37°C, 5% CO_2_ in DMEM with 10% fetal bovine serum (FBS), 1% penicillin/streptomycin, and 1% L-Glutamine. SARS-CoV-2 viral stocks were generated as previously described[10, 88, 89]. Briefly, Vero E6 cells were cultured in DMEM with 2% FBS + 10mM HEPES buffer for 1 day prior to inoculation with the SARS-CoV-2 USA-WA1/2020 strain (BEI resources, NR-52281) (GenBank MN985325.1) using a low multiplicity of infection (MOI, 0.001), in order to generate an initial viral seed stock. At 72 hpi, tissue culture supernatants were collected and clarified by centrifugation, aliquoted and stored at −80°C. The virus stock obtained from BEI Resources was a passage 4 (P4) stock and was used to generate a master seed stock (P5, or P0’) and working stock (P6, or P1’). Viral stock titers were determined by standard plaque forming assay (PFU/ml) as described below. Only stocks passaged once after seed stock (P1’) were used for experiments. All work with infectious SARS-CoV-2 was carried out in a biosafety level 3 (BSL3) facility following approved protocols.

IAV H1N1 strains A/Puerto Rico/8/1934 (PR8) and A/California/07/2009 (pH1N1) were propagated in the allantoic cavity of 11 days-old embryonated chicken eggs. At 48 hpi, allantoic fluid were collected and aliquots stored at −80°C.

### Culturing of human 3D in vitro respiratory tissue models

Human tracheobronchial air liquid interface (ALI) cultures (“Epiairway”) and human alveolar ALI cultures (“Epialveolar”) were obtained from MatTek Life Sciences (MA, USA) and cultured according to the manufacturer’s recommended protocol. Epiairway tissues were obtained at day 15 or day 21 post-seeding of primary donor lung cultures. Epiairway tissues obtained at day 15 post-seeding of primary donor lung cultures were further matured in-house in ALI interface for 7 days after receiving with 5 ml basolateral media changes every other day, with mucus washes (400 microliters 1X PBS on apical side) every 3-4 days during maturation, prior to infection. Epialveolar tissues were obtained at day 21 post seeding of primary donor lung cultures and reconstituted overnight with 5ml and 75 μl of Epialveolar media at the basal and apical sides of the tissue, respectively. Medium was changed after overnight recovery, prior to infection. Every other day, 5 ml basolateral media changes (and 75 μl apical media changes for Epialveolar tissues) were performed in both tissues for the duration of the experiments. For all SARS-CoV-2 infection kinetic studies, tissues were infected at day 23.

### Viral infection of lung tissue equivalents with SARS-CoV-2 or IAV

Tracheobronchial tissues were infected at day 21 or later of maturation to maximize matured ciliated cell populations at time of infection. Prior to viral inoculation, mucus was removed by washing twice the apical surface of tissues with 400 µl of TEER buffer (1X PBS with magnesium and calcium). Tissue inserts were inoculated with 1×10^5^ PFU of SARS-CoV-2 (for 36 h time-points and antiviral drug screening) or MOI of 0.1 and 1 of IAV (A/Puerto Rico/8/1934 or A/California/04/2009) or MOI of 0.1, 1, 3, or 10 of SARS-CoV-2 (2019-nCoV/USA-WA1/2020), assuming an average of 900,000 cells/tissue insert, for 1 h (IAV) or 1-4 h (SARS-CoV-2). Viral inoculum was removed and tissue inserts washed with PBS before continued culture. Inserts were cultured in ALI at 37°C, 5% CO_2_ for 24-36 h for antiviral compound validation or 24, 48, 72, and 144 h for viral kinetics profiling, with basal media changes every other day. Alveolar tissues were infected post day 21 with either SARS-CoV-2 or IAV in the manner described above. Multiplicities of infection were calculated based on an average of 600,000 cells/tissue insert). No pre-infection apical washes were carried out for alveolar tissues. Basal media was replaced with fresh media every other day. In alveolar cultures, 75ul of apical media was exchanged every other day. For both tracheobronchial and alveolar ALI tissues, mock-infected controls were treated in an identical fashion to viral inoculated controls.

### Plaque assay for SARS-CoV-2 production

At the corresponding time-points, secreted SARS-CoV-2 was captured by washing the apical tissue with 500 µl or 150 µl of pre-warmed tissue media (for tracheobronchial and alveolar ALI tissues, respectively). Plaque assay to determine viral loads of SARS-CoV-2-infected tissue culture supernatants was performed as previously described (Park et al., 2020; Oludanni et al., 2020). Briefly, Vero E6 monolayers in a 96-well plate format (4⍰×⍰10^4^ cells/well, performed in duplicate) were infected with 10-fold serial dilutions of collected apical supernatants in infection media (DMEM supplemented with 1% PSG). After viral adsorption (1h at 37°C, 5% CO_2_), cells were washed with PBS and incubated in post-infection media (DMEM supplemented with 2% FBS, 1% PSG) containing 1% microcrystalline cellulose (Avicel, Sigma-Aldrich) at 37°C in 5% CO_2_ for 24 h. Plates were then inactivated in 10% neutral buffered formalin (ThermoFisher Scientific) for another 24⍰h prior to removal from the BSL3. For immunostaining, fixed monolayers were washed with PBS three times, permeabilized with 0.5% Triton X-100 for 15⍰min at room temperature (RT), and blocked with 2.5% bovine serum albumin (BSA in PBS) for 1⍰h at 37°C, followed by incubation with a SARS-CoV N cross-reactive monoclonal antibody (MAb, at 1⍰µg/ml), 1C7C7, diluted in 1% BSA for 1⍰h at 37°C. After incubation with the primary MAb, cells were washed three times with PBS, and developed with the Vectastain ABC kit and DAB Peroxidase Substrate kit (Vector Laboratory, Inc., CA, USA) according to the manufacturers’ instructions. Viral counts were performed using the C.T.L. Immunospot v7.0.15.0 Professional Analysis DC and calculated as PFUs/insert.

### Focus forming unit (FFU) assay for IAV

At indicated time-points, secreted IAV was captured by washing the apical side of the tissues with 200 µl of 1X PBS. IAV titers produced from tracheobronchial and alveolar ALI tissues were measured by focus forming unit assay. Rhesus monkey kidney epithelial cells LLC-MMK2, overexpressing SIAT1 were seeded 1 day prior in black, 96-well, clear bottom plate to reach a confluency of 95-100% at time of FFU assay. Apical washes containing secreted virus from lung tissue equivalents was diluted in 2% FBS containing EMEM media and used to inoculate LLC-MMK2-SIAT1 cells for 2 h at 37°C. Viral inoculum was removed and replaced with an Avicel-media overlay and cells incubated at 37C/5%CO2 for 48 h. After 48 h, the overlay was removed, cells washed twice with 1X PBS prior to fixation with 4% paraformaldehyde. Fixed cells were washed with 1X PBS three times prior to immunostaining for IAV NP protein and counterstain with Hoechst. All plates were imaged on the InCell2200 and FFU quantified using Columbus Analysis software. All antibodies can be found in **Supplementary Table 1**.

### Drug treatments

All compounds were dissolved in DMSO unless otherwise specified. DMSO or compounds were diluted at indicated concentration directly into the basolateral media chamber of the tissue inserts for one hour prior to viral exposure and remained in the media for duration of experiment (24 h for IAV, 36 h for SARS-CoV-2). Hydroxychloroquine was dissolved in water.

### Immunofluorescence staining and analysis

Tissue inserts were completely submerged in 4% paraformaldehyde (PFA) solution for a minimum of 1.5 h or 30 min (if analyzed inside the BSL3) in 12- or 24-well plates (for EpiAlveolar and Epiairway tissues, respectively) before removal of PFA and washing with PBS three times. Tissues were permeabilized in a 0.3%-0.5% Triton X-100 in PBS solution for 15 min, followed by blocking in PBTG (1% BSA + 5% goat serum + 0.1% Triton X-100 in PBS) for 30 min to 1 h at RT or overnight at 4°C. Tissues were then stained directly in inserts or removed from inserts using a scalpel and stained whole or as cut into four equal quarters. Primary antibodies were diluted in PBTG (see **Supplementary Table 1**) and incubated at 4°C overnight or for 1h at 37°C. Secondary antibodies were also diluted in PBTG (1% BSA + 5% goat serum + 0.1% Triton X-100 in PBS) and incubated for 2 h at RT or 1h at 37°C followed by three washes with 1X PBS. Hoechst or DAPI were used to stain DNA (nuclei) of tissues infected with IAV and SARS-CoV-2, respectively. Tissues were imaged in the transwell insert or mounted in glass-bottom plates in an automated high content confocal microscope (Opera Phenix, Perkin Elmer) or using the Cytation 5 cell imaging multi-mode reader (Biotek) at 4X magnification, WFOV mode with laser autofocus; whole well images were acquired and analyzed using Gen 5 v3.8.01 software.

### qRT-PCR for quantification of SARS-CoV-2 RNA

Total RNA was isolated using TRIzol Reagent (Invitrogen) and purified using RNA Clean & Concentrator Kits (Zymo Research). 1 µg of total RNA was used to synthesize cDNA using the M-MLV Reverse Transcriptase Kit with Random Primers (Invitrogen). Gene specific primers targeting 18S RNA (forward:AACCCGTTGAACCCCATT, reverse:CCATCCAATCGGTAGTAGCG) or the SARS-CoV-2 N gene (forward: TTACAAACATTGGCCGCAAA, reverse: GCGCGACATTCCGAAGAA) and Power SYBR Green PCR Master Mix (Applied Biosystems) were used to amplify cellular RNA and viral RNA by QuantStudio 6 Flex Real-Time PCR Systems (Applied Biosystems). The relative expression levels of SARS-CoV-2 N gene was calculated using the standard curve method and normalized to 18S ribosomal RNA as an internal control.

### Cytokine and Chemokine Quantification

Basal media was collected at different time-points (24, 48, 72, and 144 hpi) from both tracheobronchial and alveolar infected ALI tissues and used to measure TH1/TH2 responses and growth factors with a customized 22-multiplex panel Human Magnetic bead Luminex assay (R&D Systems, MN, USA), following the manufacturer’s instructions. Luminex assays were performed in the BSL3 and final samples decontaminated by an overnight incubation in 1% formalin solution before readout on a Luminex 100/200 System running on Xponent v4.2, with the following parameters: gate 8000–16,500, 50⍰μl of sample volume, 50–100 events per bead, sample timeout 60⍰s, low PMT (LMX100/200: Default). Acquired data were analyzed using Millipore Sigma Belysa™ v1.0.

### Statistical analyses

Statistical significance was determined using Prism v9.0.1 software (GraphPad Software, San Diego, CA). The unpaired, two-tailed Student’s *t* test was used for two group comparisons for each time-point and reported as \**p*⍰<⍰0.05; \*\**p*⍰<⍰0.005; \*\*\**p*⍰<⍰0.0005, \*\*\*\**p*⍰<⍰0.00005.

## Conflict of Interest

There are no conflicts of interest to report.

## Authors Contribution

**EML, MF, HZ, AAG, JBT, LSM, SC, and JY** made substantial contributions to the conception, writing, and editing of this manuscript. All authors have read and agreed to the published version of the manuscript.

## Funding

Work at NCATS was funded by the by the Intramural Research Program of the National Center for Advancing Translational Sciences, National Institutes of Health (ZIA Project # TR000414-01)” to MF; by the Robert J. Kleberg and Helen C. Kleberg Foundation to JBT, by the DIR, NIAID to JY. J.B.T., A.A.G. and A.G.V. were partially supported by Robert J. Kleberg, Jr. and Helen C. Kleberg Foundation.

## Acknowledgements

We thank Jon Inglefield and Yanyu Wang at NCI/NIH and Laura Parodi and Dr. Luis Giavedoni at Texas Biomed Biology core for support in running Luminex assays. We thank Paul Shinn, Crystal McKnight, and Zina Itkin for support in compound management. We thank Matthew Hall and Richard Eastman for discussions. We thank Caroline Strong for assistance in statistical analysis. We thank the 3D Tissue Bioprinting Lab for helpful discussions and support. We thank Viktor Karetsky and MatTek for helpful discussions and support.

## Notes

### Competing Interest Statement

The authors have declared no competing interest.

