## Supplemental Figures and Tables for "Modeling SARS-CoV-2 and Influenza Infections and Antiviral Treatments in Human Lung Epithelial Tissue Equivalents"

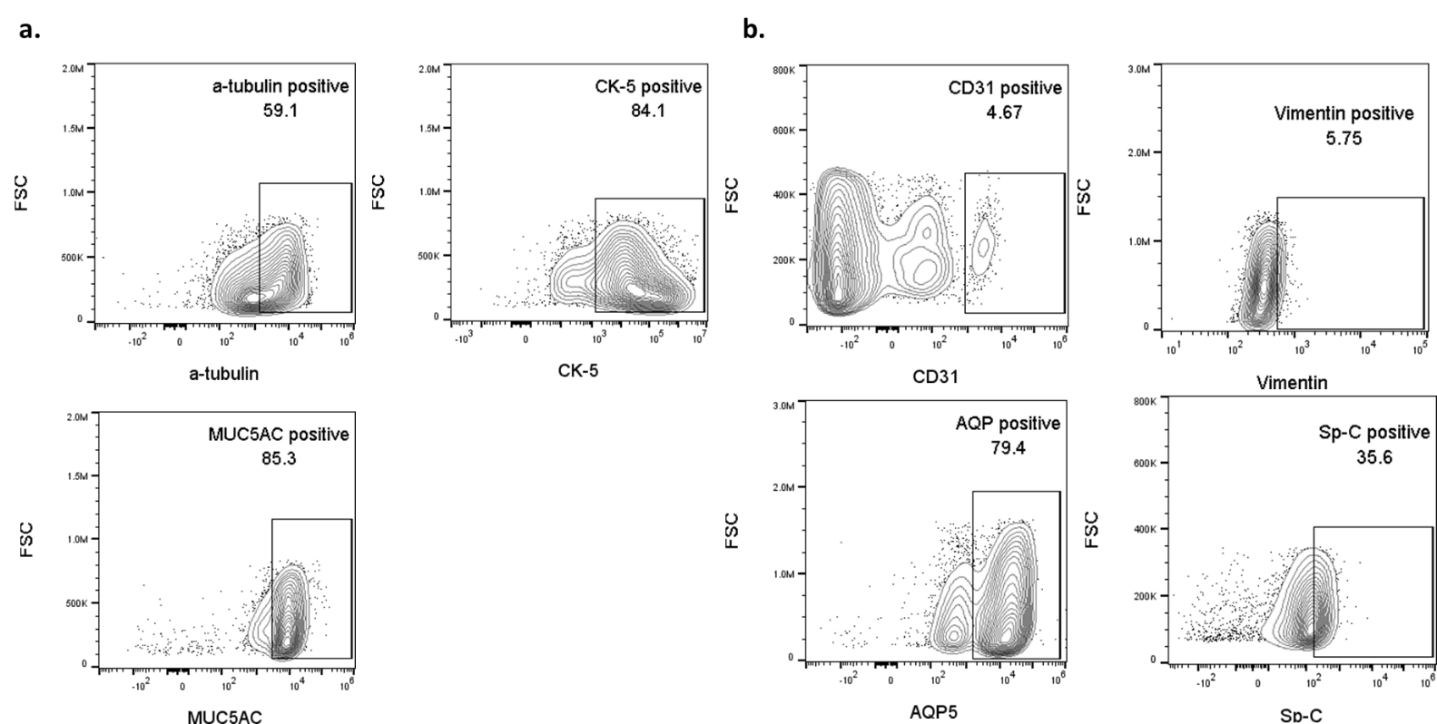

**Supplemental Figure 1: Flow cytometry analysis of alveolar or tracheobronchial tissue cultures. (a)** tracheobronchial tissues cell markers in clockwise fashion: 59.1% alpha-tubulin (ciliated cell marker), 84.1% cytokeratin-5 (basal cell marker), 85.4% MUC5AC (goblet cell marker); **(b)** alveolar tissues cell markers from left to right, top to bottom: 4.56% CD31 (endothelial cell marker), 5.22% vimentin (fibroblast marker), 79.4% aquaporin 5 (ATI/II cell marker), and 35.4% surfactant-C (ATII cell marker).

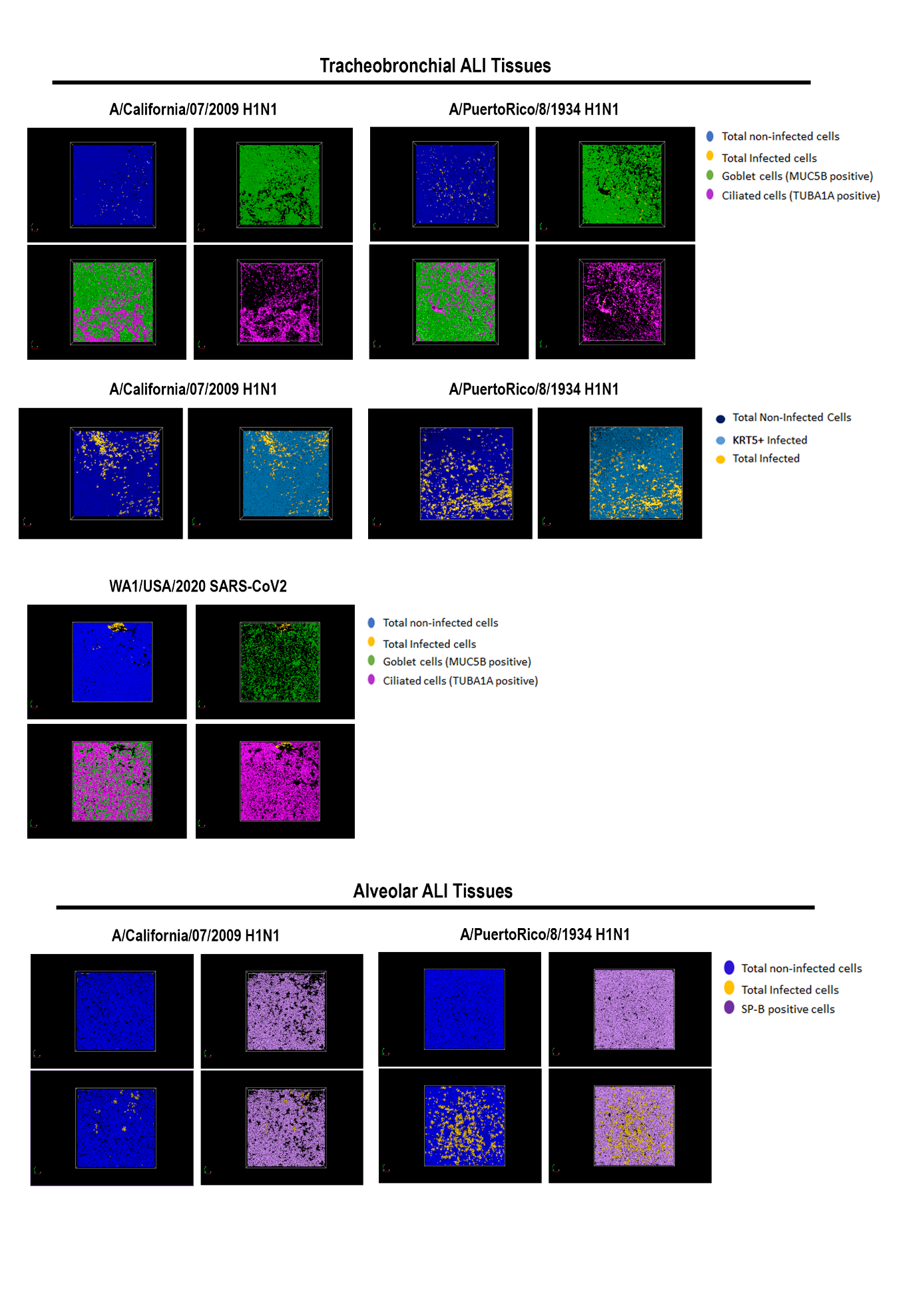

**Supplemental Figure 2: Biorender image of co-staining of cellular markers and viral antigen.** Tracheobronchial and alveolar ALI tissues were infected with IAV pH1N1 or PR8 at approx. MOI of 0.1, and SARS-CoV-2 at approx. MOI of 0.1 and fixed at 24hpi (IAV) or 36hpi (SARS-CoV-2). Representative stained images of tracheobronchial ALI tissues with Hoechst (nuclei marker, blue), α -tubulin (ciliated cell marker, magenta), MUC5B (goblet cell marker, green) and KRT5 (basal cell marker, aqua). In orange is anti-IAV NP or anti-SARS-CoV-2 spike/nucleocapsid. Bottom, alveolar: Representative stained images of alveolar ALI tissues with Hoechst (nuclei marker, blue), surfactant protein B (SP-B, ATII/pneumonocyte type II cell marker, purple) and anti-IAV NP or anti-SARS-CoV-2 spike/nucleocapsid (orange). Biorender images were created using Columbus High Content Profiler (Perkin Elmer).

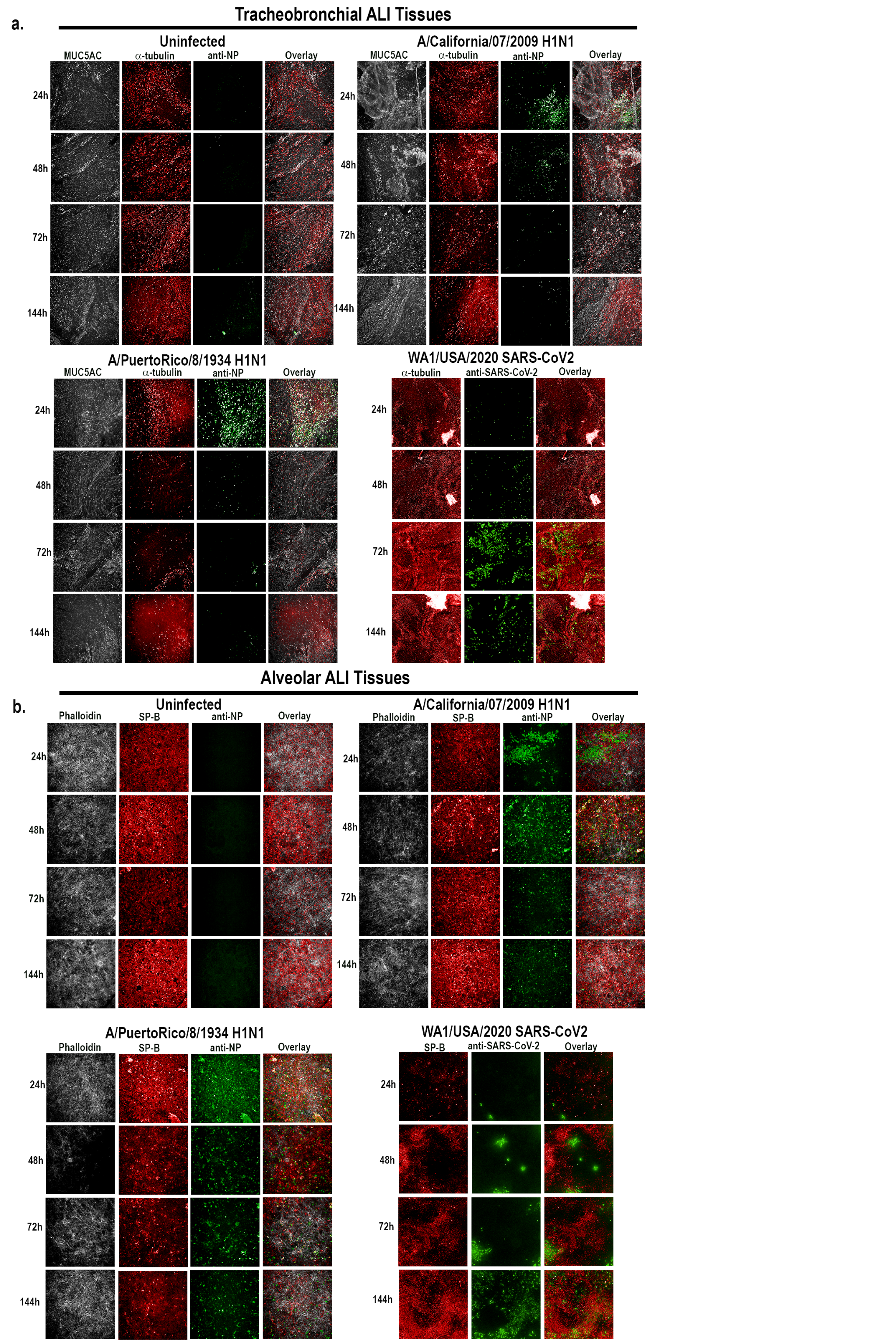

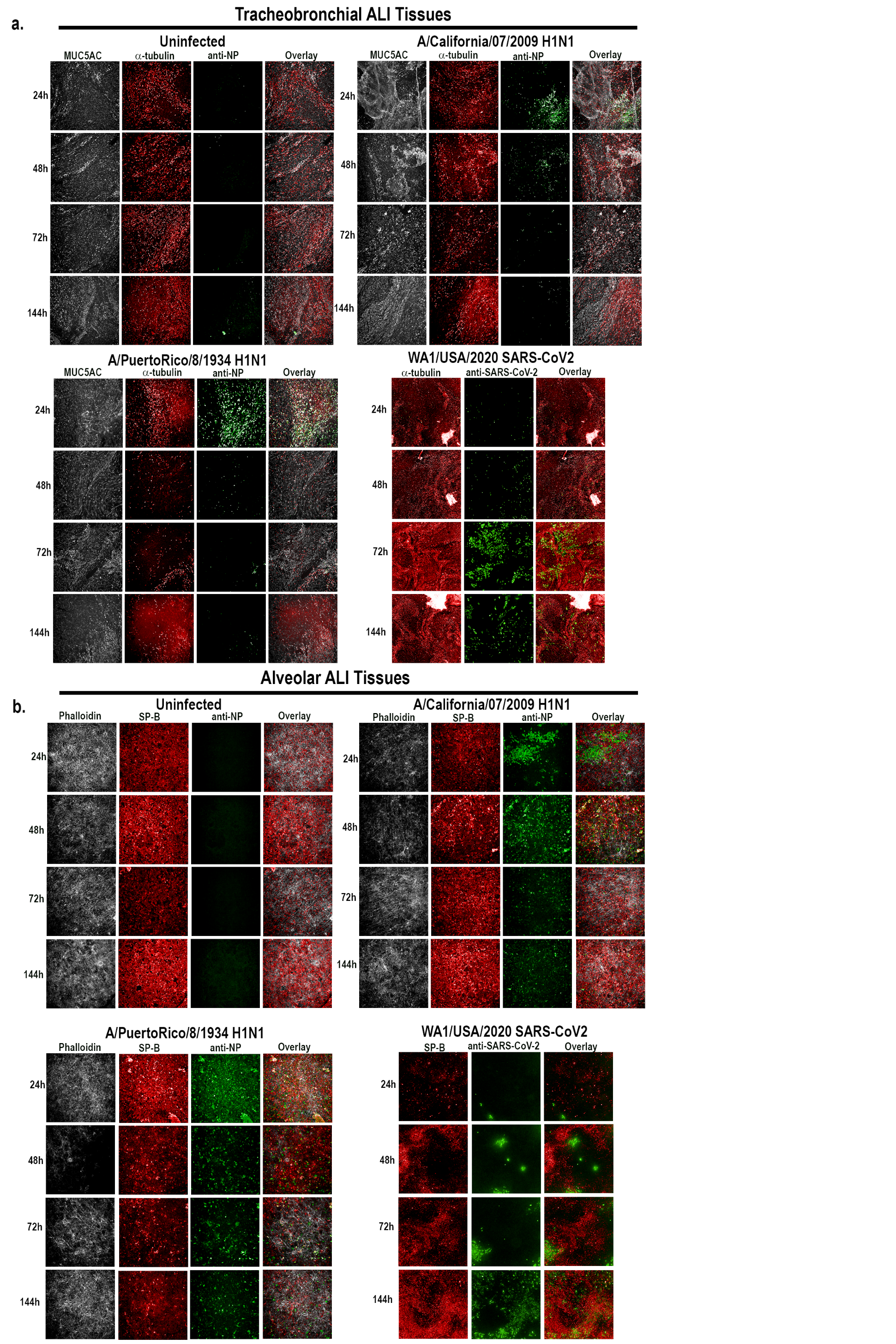

**Supplemental Figure 3: Co-staining of additional cellular markers from infection time course in Figure 2.** Tracheobronchial and alveolar ALI tissues were infected with IAV pH1N1 or PR8 at MOI of 0.1, and SARS-CoV-2 at MOI of 1 (fixed tissue samples shown) or as indicated in titer plots. Apical washes were collected and tissues fixed at 24, 48, 72 and 144 hpi**. (a)** Tracheobronchial and (**b)** alveolar ALI tissues were stained with anti-IAV NP protein and anti-SARS-CoV-2 NP/spike monoclonal antibody cocktail to label infected cells (shown in green) as well as **(a)** MUC5AC (white), alpha-tubulin (red) or **(b)** phalloidin (white), and surfactant B protein (red).

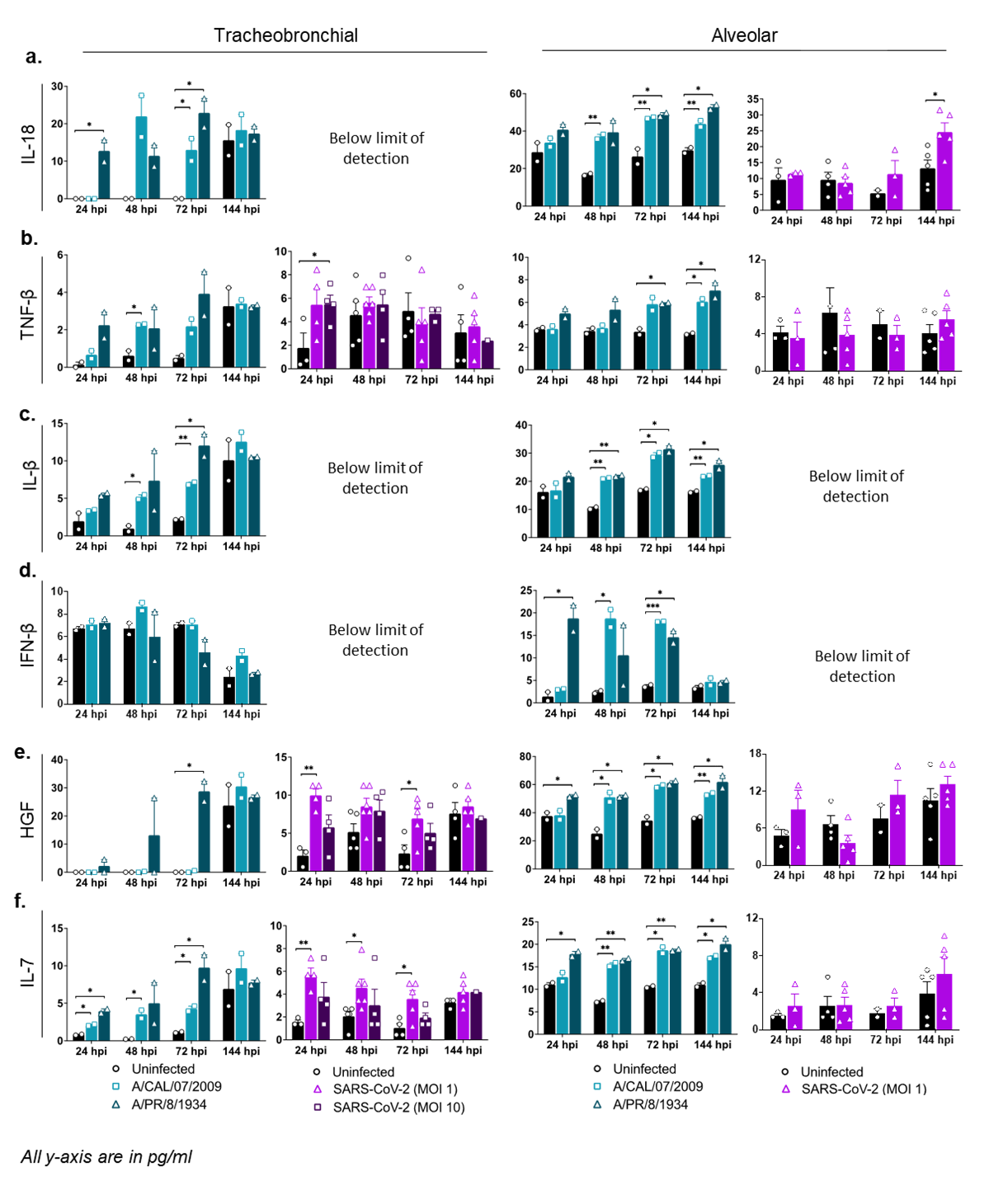

**Supplemental Figure 4: Production of other inflammatory markers.** Basal compartment media were collected from tracheobronchial (left two panels) or alveolar (right two panels) ALI tissues at indicated time-points and analyzed for cytokine and chemokine secretion by Luminex assay. IAV infected tissues (MOI of 0.1) are represented in shades of teal, where light teal shows infection with the IAV pH1N1 strain and dark teal shows infection with the IAV PR8 strain, whereas SARS-CoV-2 infected tissues are represented in shades of purple, with progressing color from low MOI (1) to high MOI (10): **(a)** IL-18, **(b)** TNF-β, **(c)** IL-1β, (**d)** IFN-β, **(e)** HGF, **(f)** IL-7. All measurements on y axis are in pg/ml. Data is represented as M±SEM for a minimum of n=2 independent experiments and/or biological replicates; Student t-test of IAV or SARS-CoV-2 infected tissues *vs*. uninfected controls at each timepoint: **p* < 0.05, ***p* < 0.005, ****p* < 0.0005, *****p* < 0.00005.

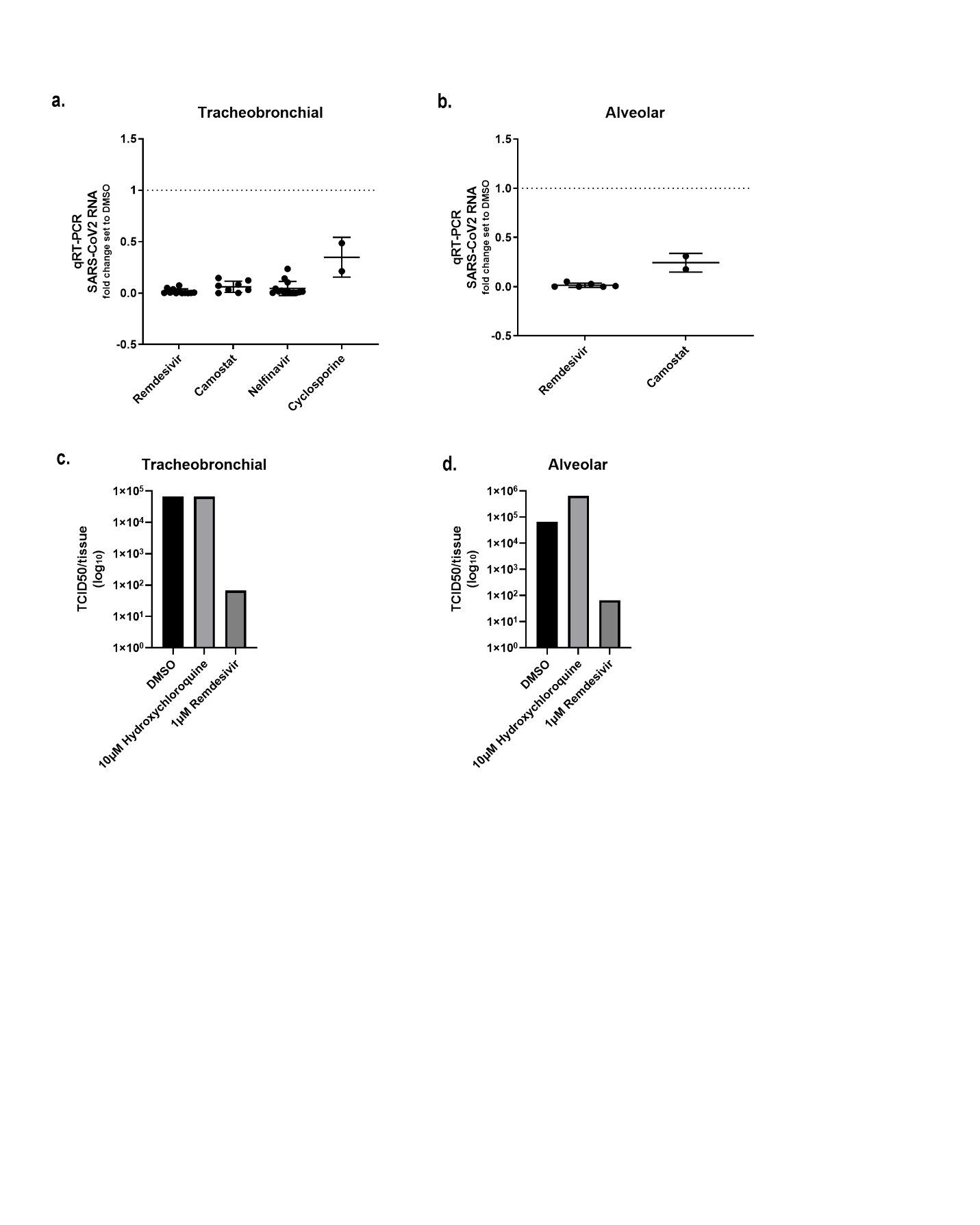

**Supplemental Figure 5: Reduction in SARS-CoV-2 viral RNA or TCID50 units in tracheobronchial or alveolar tissues treated with compounds.** **(a,b)** Intracellular SARS-CoV-2 viral RNA detection by qRT-PCR in **(a)** tracheobronchial tissues or **(b)** alveolar tissues treated with indicated compounds at 10µM at 36hpi. **(c, d)** TCID50 units measured from apical washes from (a) tracheobronchial tissues or (b) alveolar tissues treated with 10µM hydroxychloroquine or 1µM remdesivir at 48hpi.

**Supplementary Table 1:**

| **IFA/IHC Reagents** | | |
| --- | --- | --- |
| *Antibody target* | *Antibody type* | *Dilution* |
| α-tubulin (ciliated cell marker, rat mAb, ThermoFisher, MA1-80017,) | Primary | 1:200 |
| SARS-CoV-2 N (rabbit mAb, Genetex, GTX635679)  SARS-CoV-2 S (rabbit mAb, Genetex, GTX135356) | Primary | 1:100 (mixed 50/50 nucleocapsid + spike) |
| N 1C7C7 (SARS-CoV-2 marker, mouse mAb, Leinco, LT7000) | Primary | 1:500 |
| Surfactant protein B (ATII cell marker, rabbit pAb, Abcam ab40876) | Primary | 1:100 |
| AQP5+ (ATI cell marker, rabbit mAb, Abcam, ab92320) | Primary | 1:100 |
| Alexa Fluor 488 Phalloidin, ThermoFisher |  | 1:40 |
| Anti-Influenza A virus NP Mouse Monoclonal Antibody [clone: H16-L10-4R5 (HB-65), VWR] |  | 1:1000 |
| Hoechst 33342, ThermoFisher |  | 10µg/ml |
| Goat anti-Mouse IgG (H+L), Goat anti-Rat IgG (H+L), Goat anti-Rabbit (H+L) Highly Cross-Adsorbed Secondary Antibody, Alexa Fluor 488, ThermoFisher | Secondary | 1:300 |
| Goat anti-Mouse IgG (H+L), Goat anti-Rat IgG (H+L), Goat anti-Rabbit (H+L) Highly Cross-Adsorbed Secondary Antibody, Alexa Fluor 567, ThermoFisher | Secondary | 1:300 |
| Goat anti-Mouse IgG (H+L), Goat anti-Rat IgG (H+L), Goat anti-Rabbit (H+L), Alexa Fluor 647, ThermoFisher | Secondary | 1:300 |
| DAPI, ThermoFisher, 62248 |  | 1:1000 |
